# Neuromodulatory circuit effects on *Drosophila* feeding behaviour and metabolism

**DOI:** 10.1101/086413

**Authors:** Anders Eriksson, Marlena Raczkowska, Rapeechai Navawongse, Deepak Choudhury, James C. Stewart, Yi Ling Tang, Zhiping Wang, Adam Claridge-Chang

## Abstract

Animals have evolved to maintain homeostasis in a changing external environment by adapting their internal metabolism and feeding behaviour. Metabolism and behaviour are coordinated by neuromodulation; a number of the implicated neuromodulatory systems are homologous between mammals and the vinegar fly, an important neurogenetic model. We investigated whether silencing fly neuromodulatory networks would elicit coordinated changes in feeding, behavioural activity and metabolism. We employed transgenic lines that allowed us to inhibit broad cellular sets of the dopaminergic, serotonergic, octopaminergic, tyraminergic and neuropeptide F systems. The genetically-manipulated animals were assessed for changes in their overt behavioural responses and metabolism by monitoring eleven parameters: activity; climbing ability; individual feeding; group feeding; food discovery; both fed and starved respiration; fed and starved lipid content; and fed/starved body weight. The results from these 55 experiments indicate that individual neuromodulatory system effects on feeding behaviour, motor activity and metabolism are dissociated.

## Introduction

The brain is responsible for maintaining energy homeostasis by monitoring nutritional status and inducing corresponding changes in metabolism, behaviour, and food intake (Loftus et al. 2000; Leibel et al. 1995; Obici et al. 2002; Harris-Warrick & Marder 1991; Pfaff et al. 2008). Many of the brain systems that regulate metabolism and feeding are evolutionarily conserved between mammals and invertebrates, including neuromodulators (Baker & Thummel 2007; Melcher et al. 2007). Studying neuromodulators in mammals is made challenging by the expense of experiments and the brain’s immense complexity. These obstacles are overcome with the use of a major invertebrate genetic model system: the vinegar fly, *Drosophila melanogaster*. Fly neurogenetic tools enable the temporally-resolved manipulation of specific neurons, which minimizes unwanted developmental effects or compensatory mechanisms and permits the analysis of their acute role in energy homeostasis, metabolism, and feeding-related behaviours (Owusu-Ansah & Perrimon 2014; Kaun et al. 2012; Pandey & Nichols 2011).

Several neuromodulators are believed to have an effect on feeding behaviour: serotonin has been shown to regulate food intake, stomatogastric responses and appetite (Neckameyer 2010; Shimada-Niwa & Niwa 2014; Schoofs et al. 2014; Gasque et al. 2013); octopamine is believed to modulate feeding and is also regulated by the fly homologs of two obesity-linked genes (Williams et al. 2014); neuropeptide F (NPF) has been shown to respond to feeding-associated signals to help regulate feeding (Shen & Cai 2001; Lee et al. 2004); and dopamine signalling is required for normal food intake (Riemensperger et al. 2011; Zhou & Palmiter 1995)

In the present study we examined the effects of silencing neuromodulatory systems on appetitive control, food driven behaviours, metabolism and locomotion. We hypothesized that lesions in these neuromodulatory systems would have coordinated effects on behaviour, feeding and metabolism; for example, that a lesion that increased behavioural activity would also concomitantly increase metabolism and feeding. To examine this hypothesis, we used five transgenic lines to examine the circuit function of five neuromodulators (dopamine, octopamine, tyramine, serotonin, and neuropeptide F) in 11 different assays. We found that feeding phenotypes were not necessarily associated with metabolic or behavioural changes. These results indicate dissociated neuromodulator function—eroding confidence in the hypothesis—and suggesting that the organization of feeding, behaviour and metabolism requires that several modulatory systems act in a coordinated manner.

## Results

### Systematic literature review of neuromodulators and food intake in *Drosophila*

A total of 120 articles were identified in the initial search of PubMed; applying the selection criteria reduced this number to three articles (Figure1). The three identified articles contained five different experiments that utilized two different methods for assessing food intake: the CAFE assay and the proboscis extension reflex (PER) test. *Drosophila* feeding behaviour was characterized using these two assays after either silencing or activating dopaminergic, serotonergic, NPF-ergic and octopaminergic (or tyraminergic) neurons (Figure2) (Marella et al. 2012; Williams et al. 2014; Inagaki et al. 2012). The relative paucity of data relating adult *Drosophila* neuromodulator function to feeding and metabolism led us to conduct new experiments on this topic.

**Figure 1.**
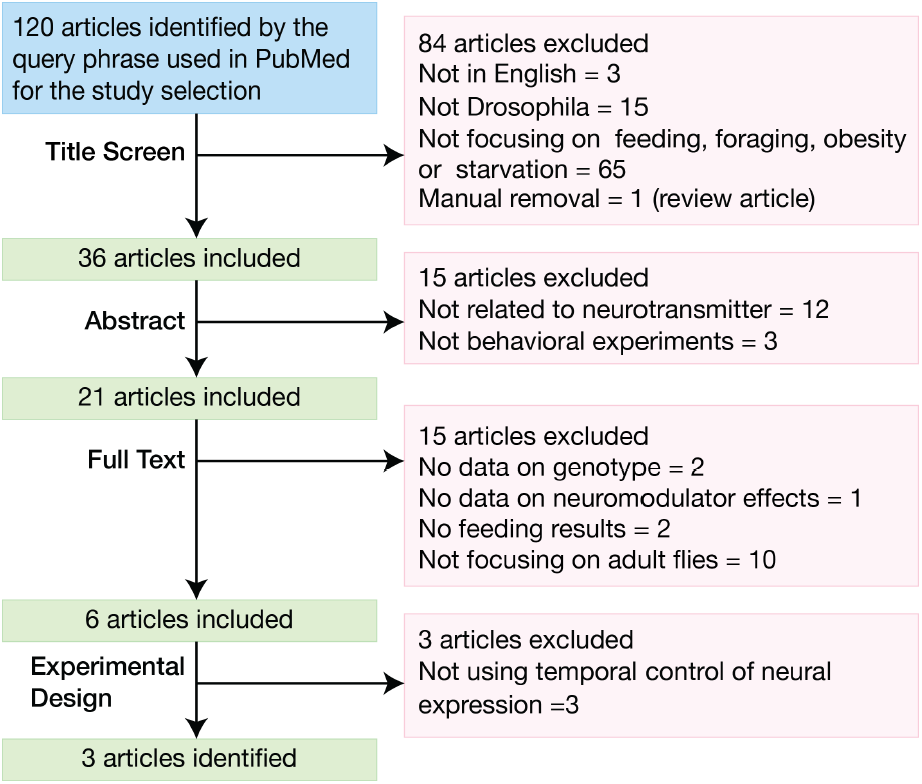
A systematic literature review of the effects of neuromodulatory neurotransmission on feeding behaviours in *Drosophila*. PubMed was interrogated using the search expression [(*Drosophila* or fruitfly or “vinegar fly” or “fruit fly”) AND (feeding or obesity or foraging or starvation) AND (NPF or “neuropeptide F”or octopamine or serotonin or dopamine or tyramine)], which yielded 120 articles. Four successive screens were then used to review the resulting literature, whereby a title screen was followed by three exclusions of increasing detail: abstract, full text and experimental design. A total of 117 article were excluded by the selection criteria and only three articles were identified for comparison with our study.

**Figure 2.**
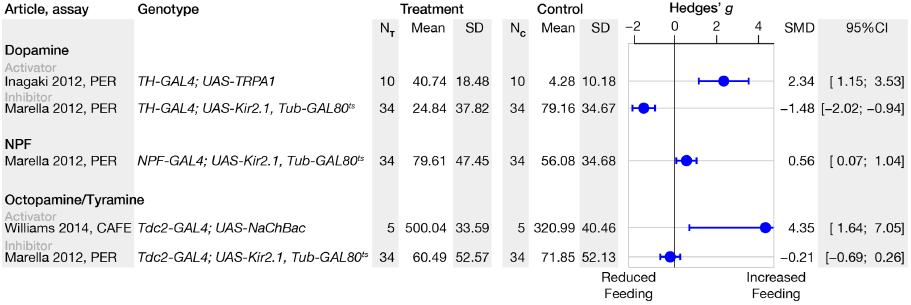
Systematic review of neuromodulators and their respective effect on feeding-related behaviors. The systematic review and the effect size is illustrated as a forest plot of standardized effect sizes (Hedges’*g*). The forest plot is grouped by neuromodulator and sub-grouped by activator and inhibitor. Error bars indicate the 95% confidence intervals (95%CI) of the standardized mean difference (SMD). Control and treatment samples sizes are given in the columns listed as NC and NT respectively. Abbreviations: *TH*, tyrosine hydroxylase; *Tdc2*, tyrosine decarboxylase 2; *NPF*, neuropeptide F precursor; CAFE, capillary feeding; PER, proboscis extension reflex; SD, standard deviation.

### Silencing of *TH-Gal4* neurons reduces activity

We used *Gal4* lines with the enhancers of five genes: *pale/Tyrosine hydroxylase* (*TH*), *Dopa decarboxylase* (*Ddc*), *Tyrosine decarboxylase 2* (Tdc2), *Tryptophan hydroxylase* (*Trh*) and *neuropeptide F* (*NPF*). These driver transgenes were used in combination with the *TubGal80*^*ts*^ conditional repressor transgene (McGuire et al. 2004) to drive expression of Kir2.1, an inward-rectifying potassium channel that silences neuronal activity (Baines et al. 1999). Following warm-treatment induction of Kir2.1 expression, motor coordination and activity were compared with two sets of control flies: a driverless line (*UAS-Kir2.1; Tub-Gal80*^*ts*^*/+*) subjected to heat, or flies maintained at 21°C that had intact Gal80^ts^ repression of *Kir2.1* expression but were otherwise genetically identical. Inhibiting electrical activity in *THGal4* dopaminergic cells had a profound effect on the activity index, defined as the proportion of time a fly spent moving. The activity index of induced *TH*>*Kir2.1* flies was reduced by –70% compared to uninduced isogenic animals (Figure 3A). Inhibiting the *Ddc-Gal4* cells, which include both dopaminergic and serotonergic neurons, reduced activity by –35%. Flies with silenced *Tdc2-Gal4* cells, which include both octopaminergic and tyraminergic neurons, exhibited a +48% increase in activity. Silencing the *Trh-Gal4* cells, which includes most of the serotonergic neurons, had only a negligible effect on activity.

**Figure 3.**
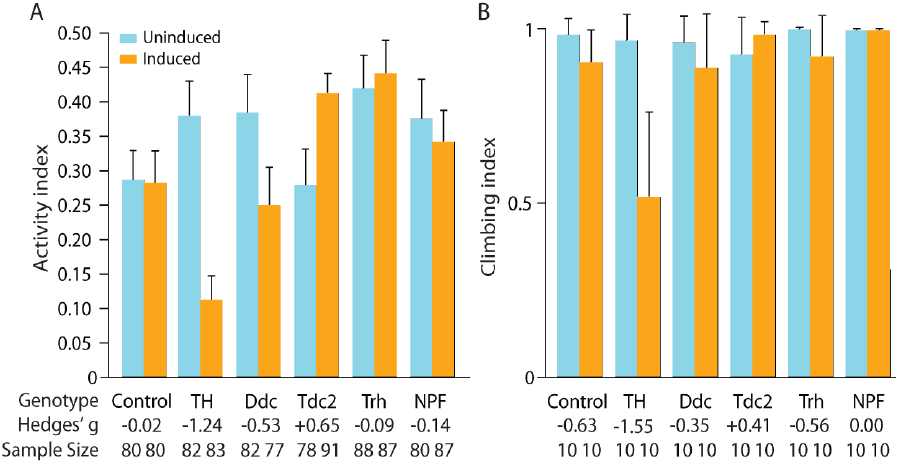
Temporal inactivation of TH-Gal4 dopaminergic neurons disrupts motor function. Activity index and climbing index for the progeny of *TH-Gal4, Ddc-Gal4, Tdc2-Gal4*, *Trh-Gal4* and *NPF-Gal4 flies crossed with Kir2.1; Tub-Gal80*^*ts*^, and a responder control line (progeny of *UAS-Kir2.1; Tub-Gal80*^*ts*^ crossed with wild type CS flies). Controls were maintained at 22 °C only (blue bars) whereas Kir2.1-induced flies were raised at 22°C before being transferred to 31°C for 24h prior to the assay (orange bars) to elicit Kir2.1 expression. The control flies carried the *UAS-Kir2.1; Tub-Gal80*^*ts*^*/+* construct but not the *Gal4* drivers. **A.** An activity index was calculated for each fly, and represented the proportion of time that the fly spent moving. *TH-Gal4*>*UAS-Kir2.1; Tub-Gal80*^*ts*^*/+* flies exhibited reduced activity: ∆activity = –0.25 [95CI –0.19, –0.32], *g* = –1.24, *P* = 1.7 × 10^-12^, N _flies_ = 82, 83. Induced *Ddc*>*Kir2.1* flies also exhibited reduced activity but to a lesser extent compared to the *TH*>*Kir2.1* flies: –0.10 [95CI –0.01, –0.18], g = –0.53, P = 0.023. Conversely, neural inhibition with *Tdc2*>*Kir2.1* resulted in increased activity: +0.67 [95CI +0.19, +0.06], g = +0.65, P = 2.10 × 10^-5^, N_flies_ = 78, 91. **B.** The climbing ability of *TH*>*Kir2.1* flies was impaired compared to controls: ∆climbing = –0.45 [95CI –0.21, –0.70], *g* = –1.55, *P* = 0.0027, N_tubes_ = 10, 10. A moderate reduction in climbing index was seen for *Trh-Gal4 flies,* ∆climbing = –0.08 [95CI –0.219, 0.001], *g* = –0.56, *P* =0.50, N_tubes_ = 10, 10. [Editor’s note: The high *P* value suggests that the sample size of this assay is too small to detect statistical significance of moderate effects.] No differences were observed in *Ddc>Kir2.1* flies. All error bars represent 95% CI. The numbers indicated below each bar denote the effect size (*g*) for the individual driver lines. The climbing index was repeated 10 times with at least five male flies, and no more than 20 flies in each vial. Abbreviations: *TH*, tyrosine hydroxylase; *Ddc*, dopa-decarboxylase; *Tdc2*, tyrosine decarboxylase 2; *Trh*, tryptophan hydroxylase; *NPF*, neuropeptide F precursor.

### Inhibition of TH cells impairs climbing ability

We next assessed climbing ability to determine whether the observed reduction in activity in the transgenic dopamine lines was associated with functional motor deficits (Bartholomew et al. 2015). Temperature-habituated flies expressing Kir2.1 in the *TH-Gal4* cells exhibited dramatically worse climbing ability compared to control flies, a climbing index reduction of –0.45. Induced *Trh*>*Kir2.1* flies displayed a moderate decrease in climbing ability, displaying a climbing index reduction of –0.08. All other induced *Gal4*>*Kir2.1* lines exhibited only small or negligible differences in climbing (Figure 3B), including a small deficit in *Ddc*>*Kir2.1*, which drives in both dopaminergic and serotonergic cells (Li et al. 2000).

### Silencing *Trh* neurons increased food intake

Neuromodulators are known to affect foraging and feeding behaviours in vertebrates and invertebrates (Shen & Cai 2001; Zhang et al. 2013; Gasque et al. 2013; Williams et al. 2014; Vucetic & Reyes 2010). Thus, we measured cumulative food intake in the five driver lines over a period of 6 h in a group of flies using a CAFE assay. Control *UAS-Kir2.1; Tub-Gal80*^*ts*^*/*+ flies drank +40% more liquid food after 31°C warm treatment, which represented the expected change in food intake as a result of warm treatment alone without Kir2.1 induction (Figure 4). Despite this large change in motor activity in response to temperature, *TH*>*Kir2.1* displayed only a minor change in food intake (Figure 4A). Interestingly, *Trh*>*Kir2.1* was the only line that displayed a substantial increase (51% over 6 h) in consumption after Kir2.1 induction and the heat recovery period (Figure 4).

**Figure 4.**
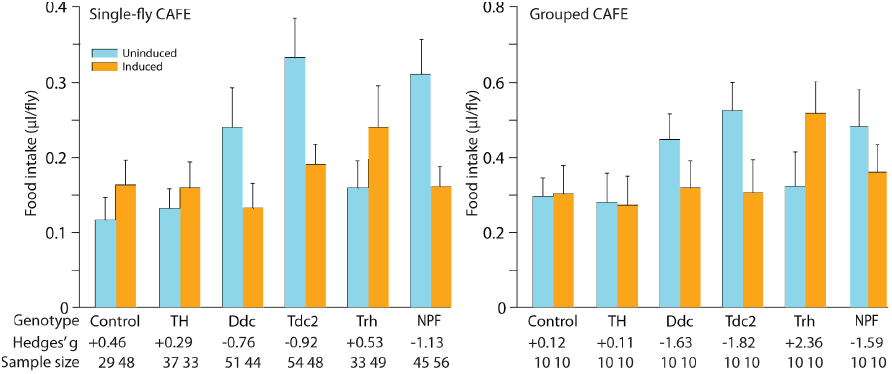
Silencing neuromodulatory circuits has effects on food intake. **A.** Flies starved for 24 hours prior to the experiment had the cumulative food intake measured over a 6 h period by capillary feeding (CAFE) assay. Control *UAS-Kir2.1; Tub-Gal80ts/+* flies drank a little more after 31°C warm treatment: Δfood intake = +0.05 μl [95CI +0.09, +0.004], *g* = +0.46 P = 0.12, N_flies_ = 29, 48. *Ddc-Gal4* flies deficient in dopamine exhibited a redu¼ction in food intake: Δ food intake = –0.11μl [95CI –0.05, –0.16], *g* = –0.76, *P* = 0.0001, n = 51, 44. Inhibition of Tdc2-Gal4 neurons also caused a substantial decrease in cumulative food consumption: Δfood intake = –0.14μl [95CI –0.10, –0.18], *g* = –0.92, *P* = 5.2 × 10^-9^, n = 54,48. Conversely, inhibition of Trh-Gal4 cells increased food consumption: Δfood intake = +0.08 μl [95CI +0.14, +0.02], *g* = +0.53, *P* = 0.052, n = 33, 49. *NPF-Gal4* flies had a reduced food intake: Δfood intake = –0.15 μl [95CI = –0.09, –0.20], *g* = –1.125, *P* = 4.65 × 10^-7^, n = 45, 56). **B.** The results of the individual CAFE assay were confirmed with conventional CAFE: groups of *Ddc-Gal4* flies had a reduced food intake with Δfood intake = –0.16 μl [95CI –0.062, – 0,27], *g =* –1.63, *P =* 0.044; *Tdc2-Gal4* Δfood intake = –0.22μl [95CI –0.15, –0.294] *g = –*1.82, *P =*0.035; *Trh-Gal4* Δfood intake = +0.19μl [95CI +0.27, +0.1], *g =* +2.36, *P =* 0.07; *NPF-Gal4* Δfood intake = –0.12μl [95CI –0.04, –0.2] *g =* –1.59, *P =* 0.04. The data represent the means with their 95% CI. A total of 10 males were used for each replicate and the assay was repeated at least five times for each genotype. The control data are depicted in blue, and the experimental (induced) data in orange. The numbers below each column denote the effect sizes for the individual driver lines. The numbers at the base of each column denote the sample size (N). Uninduced *D. melanogaster* lines exhibited increased food intake relative to the control line: *Ddc-Gal4* Δfood intake *=* +0.15 μl [95CI +20, +0.09]; Tdc2-Gal4 Δfood intake = +0.23 μl [95CI +0.29, +0.18]; NPF-Gal4 Δfood intake = +0.19 μl [95CI +025, +0.12]. Abbreviations: TH, tyrosine hydroxylase; Ddc, dopa-decarboxylase; Tdc2, tyrosine decarboxylase 2; Trh, Tryptophan hydroxylase; NPF, neuropeptide F precursor.

### Silencing *Ddc*, *Tdc2* and *NPF* cells reduced food intake

Data obtained from CAFE assays indicated that the silencing of neuronal cells in several driver lines produced decreases in feeding behaviour in flies that had been starved for 24 h. However, in all cases, the food intake of the uninduced flies was higher than that of the driverless controls: uninduced, Gal80^ts^-repressed *Ddc*>*Kir2.1*, *Tdc2*>*Kir2.1* and *NPF*>*Kir2.1* flies had +105%, +185% and +166% increased food consumption, relative to *UAS-Kir2.1; Tub-Gal80*^*ts*^*/+*controls, respectively (Figure 4A). One possible explanation is background genotypic differences between controls and experimental lines. Alternatively, since the shared genetic difference between the uninduced experimental flies and the uninduced driverless control is the presence of a driver, the increased intake could also be due to partially incomplete Gal80^ts^ repression at 22°C (Marella et al. 2012; Masek & Keene 2013; Pool et al. 2014; Jeong et al. 2016), though it is unclear why the intake effect might be an increase. Relative to the elevated baseline of uninduced *Ddc*>*Kir2.1* controls, inhibition of *Ddc* cells reduced their 6 h food intake by 0.11 μl (Figure 4A). Blockade of neural activity in *Tdc2-Gal4* cells also reduced food intake by0.14μl; similarly, silencing NPF-ergic neurotransmission decreased food intake 0.15 μl (Figure 4A). These results indicate that these neuromodulatory systems normally function to promote feeding.

### A group feeding assay confirmed two types of feeding changes

The unusual changes in feeding behaviour in the uninduced flies in the single-fly CAFE assay, and large changes in feeding in the induced flies, raised the question as to whether these data were due to isolation of the flies versus group behaviour or might be sporadic results. As such, we replicated all the experiments in a group CAFE assay (Ja et al. 2007). In the group CAFE experiment, the overall effects of genotype and induction were very similar to the original, individual-level data. The increases in the baseline feeding level were also replicated, but were less marked than in the single-fly assay: uninduced *Ddc*>*Kir2.1*, *Tdc2*>*Kir2.1, Trh*>*Kir2.1* and *NPF*>*Kir2.1* flies demonstrated a +51%, +77% and +63% increase in feeding, respectively, compared to the uninduced controls (Figure 4B). This experiment also reproduced the effect of decreased feeding after Kir2.1 induction in these lines, eliciting –29%, –42%, and – 25% reductions, respectively (Figure 4B).

### Neuromodulators influence foraging behaviour

To further assess the role of neuromodulators in feeding behaviour we assessed the effects of neural silencing on a food-discovery task (Navawongse et al. 2016) we refer to as the Small-animal Nutritional Access Chip (SNAC). Liquid food was provided to hungry flies over six sessions and the ability to approach the transient food source was measured. In each epoch, flies had 100 sec to find the food outlet alcove. Only induced *TH*>*Kir2.1* animals entered the food alcovemarkedly less than uninduced controls, with a median of 1/6 possible entries compared to 3/6: Cliff’s delta = −0.62 [95CI −0.47, −0.70], *P =* 2.5 × 10^-10^ (Figure 5A). Induced *TH*>*Kir2.1* flies also travelled a shorter distance before entering the alcove: Δ∆distance = –65 mm [-33.4, –90.7], *g* = −0.63, *P* = 2.6 × 10^-10^, n = 82, 75 (Figure 5D).Neither the time to alcove entry nor the path efficiency were affected in any of the lines, although *Tdc2*>*Kir2.1* induction resulted in a small reduction in the number of alcove entries, from a median of 3 to 2. These data suggest that neuromodulator silencing has a minimal effect on food-discovery in *D. melanogaster*, with the exception of dopaminergic neuron silencing, as observed in the induced *TH*>*Kir2.1* flies that performed poorly. This effect is likely due to the generally impaired motor function of these flies.

**Figure 5.**
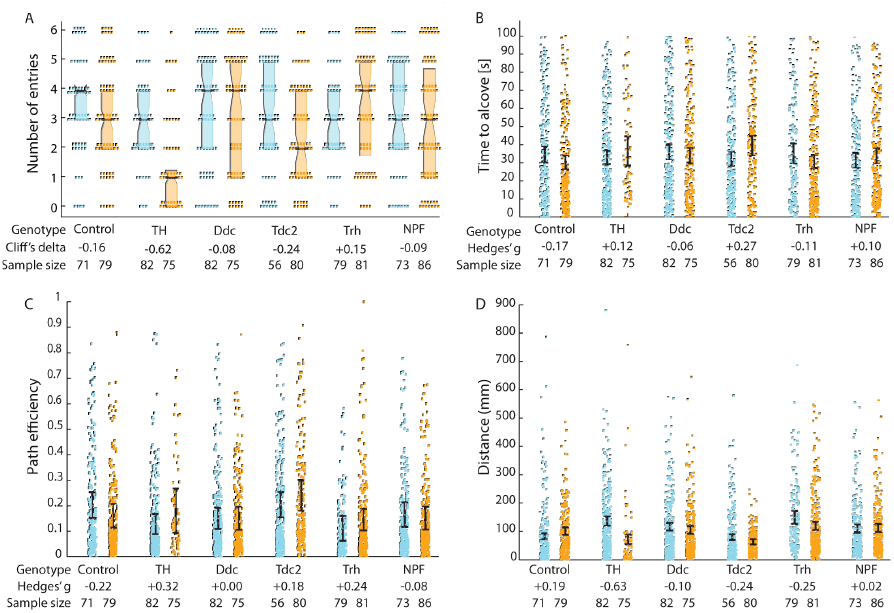
Effect of dopaminergic neuron inhibition on feeding behaviour in TH>Kir2.1 flies. **A.** Median number of entries to the feeding alcove. Induced *TH*>*Kir2.1* flies had a reduced number of entries relative to controls: Cliff’s delta = −0.62 [95CI –0.47, –0.70], *P =* 2.5 × 10^-10^n = 82, 75. *Tdc2*>*Kir2.1* reduced their number of entries to the feeding alcove, from a median of 3 to 2, Cliff’s delta Δentries = −0.24 [95CI −0.08, −0.43] = –0.8 [95CI –0.19, –1.32], *g* = –0.44, *P* = 0.006 **B.** Silencing neuromodulators did not affect the time latency to enter the alcove. **C.** No differences were seen in the path efficiency between induced and uninduced animals. **D.** Induced *TH*>*Kir2.1* flies did not travel as far to the alcove as control flies: Δdistance = –65 mm [-33.4, –90.7], *g* = –0.63, *P* = 2.6 × 10^-10^, n = 82, 75. Control flies showed no effect on distance travelled to the alcove. Flies with inhibited neurons (induced) are depicted in orange and control flies (uninduced) are depicted in blue. Dots represent the distances travelled in all epochs in which a fly successfully entered the feeding alcove. Error bars represent the 95% confidence intervals. Numbers above the columns denote the effect sizes. Numbers at the base of each scatter plot denote the sample size (N). Abbreviations: Ctrl, control; TH, tyrosine hydroxylase; Ddc, dopa-decarboxylase; Tdc2, tyrosine decarboxylase 2; Trh, Tryptophan hydroxylase; NPF, neuropeptide F precursor.

### Silencing aminergic circuits affects CO_2_ production during different nutritional paradigms

Neuromodulators are involved in metabolic homeostasis and energy regulation in both mammals and *Drosophila* (Loftus et al. 2000; Leibel et al. 1995; Obici et al. 2002; Harris-Warrick & Marder 1991; Pfaff et al. 2008). We asked, therefore, whether manipulation of neuromodulatory circuits might affect respiration in a nutritional status-dependent manner (Zhang et al. 2015). Flies were either allowed to feed *ad libitum* or were starved for 24 h prior to warm-induced Kir2.1 expression. Warm treatment of control flies at 31°_;_C caused a moderate decrease in the respiration rate of fed control flies decreased with a ΔVCO_2_ of –0.81 μl/fly/h.. Interestingly, in the starved state, silencing cells with either dopaminergic driver (*THGal4* or *Ddc-Gal4*) resulted in decreased respiration and reduced CO_2_ production where induced *TH*>*Kir2.1* and *Ddc*>*Kir2.1* decreased their CO_2_ production with 1.63 and 1.32 μl/fly/h, respectively (Figure 6B). Consistent with their increase in food intake, induced *Trh-Gal4* flies also showed an increase of 3.44 μl/fly/h in metabolism after starvation (Figure 6B).

**Figure 6.**
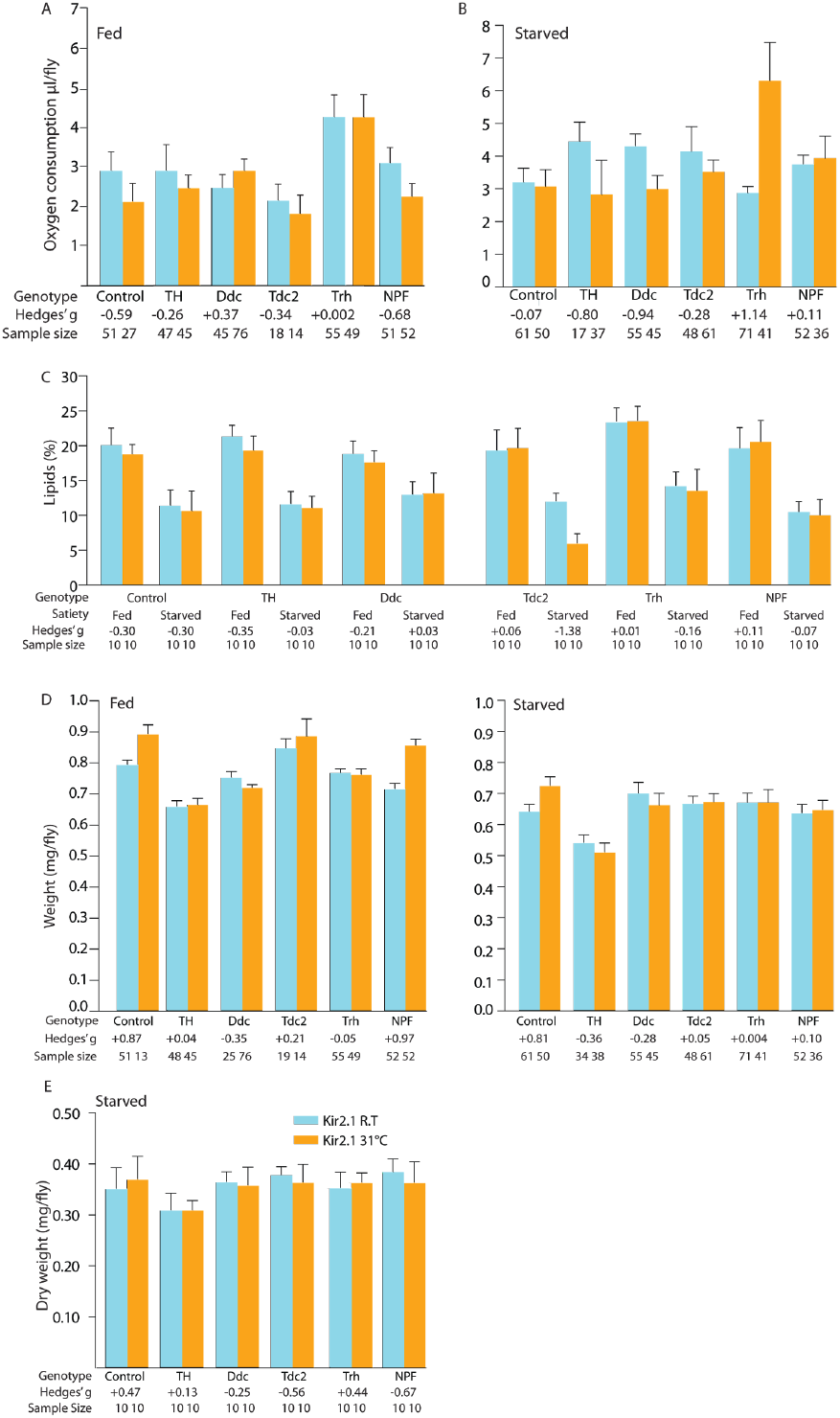
Manipulation of dopaminergic, serotonergic and NPF-ergic activity during different dietary paradigms alters metabolism. **A.** Respirometry measurements of neuromodulator driver lines with induced Kir2.1 or *UASKir2.1; Tub-Gal80*^*ts*^*/+* flies (Ctrl). The progeny were assayed in the uninduced state or after overnight warm induction to elicit Kir2.1 expression, and after *ad libitum* feeding. Several lines underwent modest changes in CO2 consumption (VCO2). Inhibition of NPF moderately decreased respiration: ΔVO2 = –0.85μl/fly/h [95CI –0.34, –1.32],*g* = –0.6835, *P* = 0.0006, n = 51, 52). Warm treatment of control flies at 31°C caused a moderate decrease in the respiration rate of fed, control flies decreased the: ΔVO2 = –0.81μl/fly/h [95CI –0.19, –1.4], *g* = –0.59, *P* = 0.0214, n = 51, 27. **B.** Respiration rate after the flies were wet starved for 24 h. Several lines underwent substantial changes in consumption: *TH*>*Kir2.1*, ΔVO2 = –1.63μl/fly/h less CO2 ([95CI –.35, –2.64], *g* = –0.6, *P* = 3.66 × 10^-5^, n = 17, 37); *Ddc*>*Kir2.1*, ΔVO2 = –1.32μl/fly/h less CO2 ([95CI –0.78, –1.83, *g* = –0.96, *P* = 2.32 × 10^-5^, n = 55, 45); *Trh>Kir2.1*, ΔVO2 +3.44μl/fly/h [95CI +4.7, +2.4], *g* = +1.14, *P* = 1.94 × 10^-10^, n = 71, 41. **C.** Whole-body lipid levels were determined in fed flies (0 h) and flies starved for 24 h. All lines underwent substantial changes in lipid levels during starvation, but these were comparable to control flies, with the exception of *Tdc2*>*Kir2.1,* which displayed a –50% loss of fat after 24 h starvation when compared to its uninduced control: Δlipid = –6.08% [95CI –4.32, –7.7], *g = –*1.38*, P* = 0.009. When comparing warm-treated wild type control flies (*UASKir2.1; Tub-Gal80*^*ts*^/+) the Δlipid of the *Tdc-Kir2.1* flies was –4.76% ([95CI –3.24, –6,92], *g* = –2.65, *P* = 0.04202) after 24 h of starvation. Five males were used for each replicate, and the assay was repeated at least five times for each genotype. **D and E.** Wet and dry whole-body weight for controls or respective genotypes in 5–7 dayold flies. Flies with impaired Tdc2 neuromodulation had an increase in body weight, No marked changes in weight were observed in any of the lines 24 h after induction, in either fed or starved animals. Flies with inhibited neurons (induced) are depicted in orange and control flies (uninduced) are depicted in blue. Data represent the means with their 95% CI; numbers at the base of each column denote the sample size (n) and the numbers at the top denote the respective effect size. Abbreviations: Ctrl, control; TH, tyrosine hydroxylase; Ddc, dopa-decarboxylase; Tdc2, tyrosine decarboxylase 2; Trh, Tryptophan hydroxylase; NPF, neuropeptide F precursor.

### Silencing neuromodulator cells elicited at most modest changes in fat and weight

We next assessed whether neuromodulator disruptions caused changes in body weight by measuring the mass and lipid content of fed and starved flies. Group comparisons were made with and without Kir2.1 induction in different sets of flies (Figure 6C). Only minor differences in lipid levels were observed in the majority of the experimental lines, in both the fed and 24 h-starved states (Figure 6). An exception was observed in the induced *Tdc2-Kir2.1* flies, which had a lipid content reduction of 6.08% compared to uninduced controls after 24 h starvation(Figure 6C); when compared with the warm-treated wild type control flies (*UAS-Kir2.1; Tub-Gal80*^*ts*^/+) the lipid content of *Tdc-Kir2.1* flies was reduced by 4.76% (Figure 6C). Interestingly, none of the flies showed any substantial changes in overall wet weight under any of the experimental conditions except for NPF flies which displayed a modest increase of +0.14 mg [0.082, 0.19], g = +0.97, *P* = 2.45 × 10^-6^ in fed flies (Figure 6D). However, control flies (*UAS-Kir2.1; Tub-Gal80*^*ts*^/+) also underwent a modest weight increase (+0.9 mg), suggesting that the weight gain attributable to NPF-ergic silencing (as opposed to a heat effect) was even more modest. Moreover, weight changes in NPF>Kir2.1 were seen neither in starved flies (Figure 6D), nor in dry weight measurements (Figure 6E). We conclude that none of the neuromodulators silenced are capable of modulating major weight changes in a 24 h interval, possibly because insufficient energy is expended over this time.

## Discussion

### Dopaminergic effects on motor function and feeding

Previous studies have shown that activating *TH-Gal4* cells in transgenic flies promotes the PER (Inagaki et al. 2012), whereas inhibiting these cells reduces the PER (Marella et al. 2012), suggesting that dopaminergic activity modulates feeding drive (Figure 2). However, our data indicate that inhibiting *TH-Gal4* cells has only a trivial effect on post-starvation food intake (Figure 4). Rather, silencing the *THGal4* cells resulted in substantial effects on overall activity, climbing, food discovery and respiration—outcomes that are consistent with a disruption in motor function (Table 1). From these data we conclude that all the phenotypes we observed in this transgenic line are due to this generalized motor deficit. These findings confirm previous work that has shown that loss of either dopaminergic neurons or the dopamine synthetic enzyme, tyrosine hydroxylase, results in motor impairments (Riemensperger et al. 2013; Riemensperger et al. 2011; Shaltiel-Karyo et al. 2012; Islam et al. 2012; Coulom & Birman 2004). Silencing the dopaminergic cells in the *Ddc-Gal4* driver did not result in motor deficits; rather, inhibiting the *Ddc-Gal4* cells (which include both dopaminergic or serotonergic neurons) robustly and specifically suppressed food intake and reduced respiration. While *Ddc-Gal4* targets some serotonergic cells, silencing cells using the nearly comprehensive *Trh-Gal4* serotonergic driver, had the opposite effect: induced *Trh*>*Kir2.1* flies increased their food intake and produced more CO_2_ (Table 1). These distinct results suggest that *Ddc-Gal4* contains a subset of dopaminergic cells (or maybe serotoninergic cells) that normally act to promote feeding. One candidate group of feeding-promoting cells is the paired antero-medial cells, which are known to promote appetitive memory (Burke et al. 2012; Liu et al. 2012) and are numerous in *Ddc-Gal4* cells but sparse in *TH-Gal4* cells.

**Table 1.** Summary of results from the behavioural and metabolic assays. Standardized effect sizes (Hedges’ *g* and Cliff’s delta) are shown for all assays conducted on neuromodulator-silenced flies. Cliff’s delta values are marked with ‘Δ’. Effect sizes indicate the difference between uninduced control flies and warm-treated flies with de-repressed Kir2.1 expression. Effect sizes are listed for all 14 metrics from 11 assays conducted on five different neuromodulator driver lines. Moderate (*g* > 0.50 or Cliff’s Δ >0.47) and larger effect sizes are coloured in red for decreases and green for increases. Silencing *TH-Gal4* cells produced coordinated decreases in motor-related phenotypes (activity, climbing, alcove entries, distance travelled). The other four lines had reproducible effects on post-starvation food intake, but these were not accompanied by consistent behavioural or metabolic phenotypes.

| Assay | *TH-Gal4* | *Ddc-Gal4* | *Tdc2-Gal4* | *Trh-Gal4* | *NPF-Gal4* |
| --- | --- | --- | --- | --- | --- |
| Activity | -1.24 | -0.53 | +0.65 | -0.09 | -0.14 |
| Climbing | -1.55 | -0.35 | +0.41 | -0.56 | -0.00 |
| Food intake - Single | +0.29 | -0.76 | -0.90 | +0.53 | -1.13 |
| Food intake - Group | +0.11 | -1.63 | -1.82 | +2.36 | -1.59 |
| Alcove entries | Δ-0.62 | Δ-0.08 | Δ-0.24 | $\Delta$+0.15 | Δ-0.09 |
| Time to alcove | +0.12 | -0.06 | +0.27 | -0.11 | +0.10 |
| Path efficiency | +0.32 | +0.00 | +0.18 | +0.24 | -0.08 |
| Distance travelled | -0.55 | -0.10 | -0.24 | -0.25 | +0.02 |
| VO2 - fed | -0.26 | +0.37 | -0.34 | +0.00 | -0.68 |
| VO2 - starved | -0.80 | -0.94 | -0.28 | +1.14 | +0.11 |
| Lipid - fed | -0.35 | -0.21 | +0.06 | +0.01 | +0.11 |
| Lipid - starved | -0.03 | +0.03 | -1.38 | -0.16 | -0.07 |
| Weight - fed | +0.04 | -0.35 | +0.21 | -0.05 | +0.97 |
| Weight - starved | -0.36 | -0.28 | +0.05 | +0.00 | +0.10 |

### Reduced feeding in animals with silenced *Tdc2-Gal4* cells

Tyrosine decarboxylase is required for the synthesis of both octopamine and tyramine (Roeder 2005). Octopamine is known to have roles in a range of behaviours (Crocker & Sehgal 2008; Crocker et al. 2010; Zhou & Rao 2008), including appetitive learning (Burke et al. 2012), and starvation-induced hyperactivity (Yang et al. 2015); less is known about the role of tyramine (Roeder 2005). Previous work found that activating the *Tdc2-Gal4* cells produces a dramatic increase in capillary feeding, while silencing these cells causes only a minor reduction in PER (Williams et al. 2014; Marella et al. 2012). In our experiments, silencing *Tdc2-Gal4* cells caused a substantial decrease in food intake, consistent with the earlier evidence that *Tdc2-Gal4* cells function to promote feeding. Additionally, inhibition of the *Tdc2-Gal4* cells was found to increase lipid depletion during starvation (Figure 6C), but it is unlikely that this response directly relates to the food intake phenotype as both parameters were measured after 24h starvation.

Other studies have identified specific neuronal populations to be involved in food intake in *Drosophila*, for example cells captured by the *c673a-Gal4* driver (Al-Anzi et al. 2009). The *c673a-Gal4* subset is a heterogeneous population containing insulin-releasing, dopaminergic and serotonergic neurons, dispersed throughout the brain. Silencing of c673a-Gal4 neurons caused an increase in food intake and fat storage, while the opposite (extreme leanness) was observed when this neuronal population was hyperactivated. These results are contrary to what was observed in the current study, but unlike the study by AlAnzi et al. our driver lines are specific for each neuromodulator while c673a-Gal4 is a heterogenous population (Al-Anzi et al. 2009). As we found that silencing of dopaminergic neurons to decrease food intake and an increase in food consumption when silencing serotonin, it is likely that the observed phenotype for c673a-Gal4 is mediated by the serotonergic population.

### Are feeding, activity and metabolism dissociable?

We originally hypothesized that perturbed neuromodulatory function would have consistent behavioural and metabolic effects. A previous study identified a broad, population of insulin-releasing, dopaminergic and serotonergic neurons that caused an increase in food intake and fat storage when silenced, and fat loss when the neuronal population was hyperactivated (Al-Anzi et al. 2009); our data suggest that the coordinated effects may be due to this population’s heterogeneous, complex nature. Both the individual and group assays of post-starvation feeding found that silencing the *Trh-Gal4* cells led to an increase in food intake (Table 1). These animals also displayed an increased respiration rate, but this effect was not accompanied by an increase in activity. Silencing *NPF-Gal4* cells produced decreases in post-starvation feeding as well as a decrease in metabolism during fed state. In line with the metabolic data, we were also able to observe an increase in wet body weight for the *NPF-Gal4 flies*, while no difference was seen in dry weight, despite having a reduced food intake. These changes indicate that *NPF*-silencing affects metabolism. Consistent with previous studies, our results are in agreement regarding *NPF* and its effects on food intake. Loss of function of *NPF* in *Drosophila* larvae has been shown to have a negative effect on food intake as it decreases appetite while overexpression increases food intake producing bigger and heavier flies (Lee et al. 2004). Even though silencing *TH-Gal4* cells elicited coordinated reductions in activity, climbing ability and CO_2_ production, these effects did not translate into reductions in feeding, something that might be expected in an animal with reduced energy expenditure from activity. These isolated phenotypes suggest that incapacitating one neuromodulatory system may have specific, dissociated effects on one or several as-pects of feeding, activity and/or metabolism. While it is possible that the diverse effects could have arisen from experimental heterogeneity or sampling error, our findings erode confidence in coordinated neuromodulation of behaviour and metabolism, our original hypothesis. Rather, the data suggest that neuromodulatory subsystems regulate physiological functions in a distinct, separable manner.

The present study is the fourth study in adult *Drosophila* to specifically examine the effects of silencing specific neuromodulatory circuits (Marella et al. 2012; Inagaki et al. 2012; Williams et al. 2014). Recently developed, specific drivers, such as split-Gal4 lines targeting dopaminergic cells (Aso et al. 2014), will allow for new analyses of subsets of the neuromodulatory systems examined here. Such future studies will help determine the extent to which multi-phenotypic effects are due to pleiotropy of single neuromodulator cell types, and which effects are due to dissociable functions within the overall system.

## Methods and materials

### Fly stocks and maintenance

The five Gal4 driver lines used were *TH-Gal4 (Friggi-Grelin et al. 2003)*, *Ddc-Gal4 (Li et al. 2000)*, *Tdc2-Gal4 (Cole et al. 2005)*, *NPF-Gal4 (Wu et al. 2003)*, *Trh-Gal4 (Alekseyenko et al. 2010)*. Driver lines were outcrossed with the Canton-Special (CS) wild type strain for five generations. Driver lines and CS were then crossed with *UASKir2.1; Tub-Gal80*^*ts*^ flies to produce the male F1 flies used in all behavioural experiments. The CS flies crossed with *UAS-Kir2.1; TubGal80*^*ts*^ were used as a negative control for each assay and are hereafter referred to as controls. Flies (5–8 days old) were maintained at 22°C, in 60-70% relative humidity, under 12:12 hour artificial light-dark cycles before the experiment day. Neuronal silencing mediated by over-expression of the potassium Kir2.1 channel was induced in flies carrying a Gal4 driver transgene and *UAS-Kir2.1; Tub-Gal80*^*ts*^ by incubation at 31°C for 24 h followed by 22°C for 24 h, 1 day prior to the start of the experiment. Post-heat recovery has been previously used with *UAS-Kir2.1; Tub-Gal80*^*ts*^, for various intervals at different temperatures: behavioral results with Gal80^ts^ control (Shih et al. 2015; Kim et al. 2012; Mann et al. 2013) and with GeneSwitch control (Osterwalder et al. 2001) of *UAS-Kir2.1* (Depetris-Chauvin et al. 2011) indicate that Kir2.1 inhibition persists for at least 24 h (Mann et al. 2013), and as long as 72 h (Depetris-Chauvin et al. 2011), after the removal of induction conditions. Note that, as electrophysiological recordings (Baines et al. 2001) from the neuromodulatory neurons were not possible, the presence of sufficient hyperpolarization to suppress action potentials in any (or all) cells cannot be affirmed. For experiments involving starvation, flies were wet starved for 24 h before the start of the experiment during the heat recovery phase; during wet starvation, flies were deprived of food but not water in a vial containing a strip of filter paper soaked in deionized water. Different sets of flies were used for the uninduced and the induced Kir2.1 groups, as well as for the fed and the food-deprived flies.

### Systematic review - database search

A systematic literature search was conducted as previously described (Yildizoglu et al. 2015). The search expression [(*Drosophila* or fruitfly or “vinegar fly” or “fruit fly”) AND (feeding or obesity or foraging or starvation) AND (NPF or “neuropeptide F ” or octopamine or serotonin or dopamine or tyramine)] was used to query PubMed in September 2016, which resulted in 120 records (Figure 1). All the bibliographic information, including Title and Abstract, was exported to a Microsoft Excel spreadsheet, and the titles, abstracts and/or full texts were interrogated. The spreadsheet was used to record the results of the full text screen, which included a detailed screen of experimental design.

### Systematic review - study selection

The literature selection process was designed to identify experiments that examined the involvement of neuromodulation on feeding in *Drosophila* by using various feeding assays. Our analysis focused on experiments that aimed to understand the role of neuromodulators on feeding by using inducible neuronal activation or deactivation in genetically modified flies. The 120 records yielded from the PubMed search were screened in four stages based on title review, abstract reading, full text scan and a detailed review of experimental design, to systematically exclude studies that were not relevant. After systematic review, 117 studies were eliminated (Figure 1), and data were collected from the remaining three studies, as detailed below.

### Systematic review - data extraction

The following data were collected from the included studies: authors, year of publication, figure and panel numbers, genotype, mean number of flies in the control and experimental groups and the corresponding standard errors of the mean and sample sizes of each group, age and gender of the flies, type of food used during feeding experiments, and the number of hours of starvation prior to starting the experiment. Numeric data were digitized from graphics with the Measuring Tool in Adobe Acrobat Pro.

### Small-animal Nutritional Access Control (SNAC) chip food discovery assay

A microfluidic feeding assay was performed as previously described (Navawongse et al. 2016). Briefly, each microfluidic SNAC chip contained a 20 × 22 mm arena with a feeding alcove connected to a microfluidic channel that is designed to deliver 70 nl of liquid food via an actuator pump. Eight chips were run simultaneously. Experiments were conducted inside an incubator maintained at 22°C. The flies were given 10 min to habituate to the chip/incubator environment prior to the start of the experiment. A combined stimulus of blue light with an 85 dB, 300 Hz sound was administered at the onset of the 70 nl liquid food reward (Navawongse et al. 2016). The liquid food (5% sucrose 10% red food dye in deionized water) was retracted automatically when the fly was detected inside the feeding alcove or a timeout of 100 sec was reached. Six foraging-feeding epochs were imposed in succession with a 120 sec inter-epoch interval. The following parameters were recorded with custom LabView code as previously described: the number of entries into the feeding alcove, time to feed, path efficiency, and the distance travelled during the time the food was presented (Navawongse et al. 2016). Task performance was calculated by dividing the number of epochs with successful alcove entries by the total number of epochs. We “time to feed” as the latency to enter the feeding alcove, and “path efficiency” as the distance of the most direct path to the feeding alcove divided by the actual distance travelled by the fly in each feeding epoch. For the ac-tivity data, two color cameras (A601fc, Basler, Germany) were used in conjunction with custom LabVIEW software to record animal motion. With custom MATLAB software, the activity index was then calculated as the proportion of time that an animal spent moving. This calculation used 0.5 s bins; in this time interval, the fly was determined to be immobile if it moved less than 1 mm. A fly was excluded for the epoch if its total distance traveled was less than 200 mm or if the activity index was below 10%. As the number of alcove entries are ordinal data, we plotted the median and reported a non-parametric effect size, Cliff’s delta. Cliff’s delta values were calculated using the orddom package (Rogman 2013) in R. While *P* values from the Mann-Whitney U distribution were reported *pro forma,* no significance tests were performed.

### Climbing assay

Male flies were separated from the F1 progeny and maintained in a freshly prepared food vial 24 h before the start of the experiment. No more than 20 flies were housed within the same vial. Five male flies were put inside a 50 ml disposable polystyrene serological pipet (Fisher Scientific, Waltham, USA) that was cut to 50mm in length. The top and bottom of the tube were sealed with parafilm that was punctured with three small holes to provide ventilation. The tube was then placed flat on a surface for 1 h at 22°C while the flies acclimatised. At the beginning of the experiment the tube was tapped 2–3 times to encourage the flies to the bottom of the tube. The time for each fly to reach the top marked point of the tube was then recorded. Any flies that could not reach the top mark within 60 sec were denoted as failing to climb. The climbing index was calculated as a range between 0 and 1 for the number of flies that managed to reach the top mark within a certain time. If they failed they got a score of 0, and a score of 1 if they succeeded. An average for all flies was calculated for all the trials.

### CO_2_ metabolism assay

Metabolic changes were analyzed in flies by measuring the CO_2_ absorbance in two sets of experimental flies: control and food deprived. Male flies were transferred to either a vial containing food (control group) or a vial with water soaked filter paper (starvation group) 24 h before the start of the experiment. Kir2.1 heat induced flies were subjected to the heat recovery period before the starvation period. All flies were maintained at 22°C under a 12:12 hour light-dark cycle. The respiration chamber consisted of a 1 ml syringe (Becton Dickinson, USA) and a glass microcapillary tube (53432-728, VWR, USA) glued to the 16-gauge needle (Becton Dickinson, USA). A thin layer of absorbent and non-absorbent cotton filled the top of syringe. A fly was anesthetized by cooling and loaded into the syringe. The syringe plunger was inserted to close the chamber, leaving a 1–1.5 cm space for the fly to move. A total of 15 μl potassium hydroxide (KOH, Sig-ma, USA) was loaded on the top of the syringe and the glass micro-capillary tube was assembled onto the syringe. The gap between the syringe and holder were wrapped with parafilm. The respiration chambers were incubated under water inside the 22°C incubator (Sanyo, Japan) for 10 min. A small volume of 30% sucrose solution containing 10% food colour dye filled the top of the glass capillary to isolate the respiration chamber from the outside environment. The position of the coloured solution was measured 15 min after it was filled and its final position was determined after 1h. The CO_2_ metabolic volume consumption during a 1 h period was obtained from the increase in volume within the glass capillary.

### Body weight and lipid measurements

Male flies were starved for 1 h under non-dehydrating conditions to limit the amount of food-derived lipids present in the gut. Each group of flies were then weighed to 0.1 mg accuracy using an analytical balance (Sartorious, Singapore). For lipid measurements, groups of five males were weighed to obtain their wet mass and then dehydrated at 65°C for 24 h and subsequently weighed to obtain their dry mass. Lipid extraction was performed by placing intact, dehydrated flies in a glass vial containing 9 ml diethyl ether for 24 h at room temperature under gentle agitation. After incubation the diethyl ether was removed and the flies were air dried for an additional 24 h at room temperature. The weight of the flies was then remeasured to obtain lean dry mass. The total lipid content of the flies was considered to be the difference between the dry mass and lean dry mass.

### Capillary feeding (CAFE) assays

Male flies were anesthetized by cooling and placed in chambers for the CAFE assay, where capillaries delivered liquid food (5% sucrose 10% red colour food dye in deionized water) to the fly. The experiment was conducted within an incubator that was maintained at 22°C (Ja et al. 2007). The individual Gal4 lines were tested on different days; all experimental conditions were kept constant for all of the driver lines in between and during each experiment with respect to: temperature, humidity, circadian time, days after eclosion and the sex of the flies. The level of the fluid was noted at the beginning of the experiment and 6 h later. A control experiment, whereby no flies were housed inside the chamber, was also conducted in order to calculate the loss of fluid due to evaporation. The difference in the reduction of fluid level between the experimental and control chambers was assigned as the volume partaken by an individual fly. We used a group assay in which 10 flies were assayed in a vial with a single capillary as previously described (Ja et al. 2007), as well as a single-fly assay, in which each capillary was accessible by a single fly kept in a 12 mm × 12 mm × 2 mm chamber cut from acrylic. The food intake assays for the individual fly lines were not performed concurrently but with identical duration, start time for the experiment, age and sex of flies, temperature and starvation time.

### Data analysis

Data were analyzed with custom scripts written in LabView, Matlab and Python, and visualized with GraphPad. The data were analyzed with estimation methods to calculate median, mean, mean differences (Claridge-Chang & Assam 2016), Hedges’ *g* and Cliff’s delta where appropriate. Standardized effect sizes for Hedges’ *g* are referred to as ‘trivial’ or ‘negligible’ (*g* < 0.2), ‘small’ (0.2 < *g* < 0.5), ‘moderate’ (0.5 < *g* < 0.8) or ‘large’ (*g* > 0.8) as per standard practice (Cumming 2012). Similarly, the Cliff’s delta uses three different thresholds: <0.147 as“negligible”, <0.33 “small” <0.47 “medium” and >0.47 as “large” for description purposes (Cliff 1993). Both Hedges’ g and Cliff’s delta are standardized measures of effect size where the later is used for ordinal and non-parametric data. Hedges’ g is an indication of how much two groups differ with each other, i.e., a *g* of 1 shows that the two groups differ by 1 standard deviation. Cliff’s delta measures how often values in one group are higher than the values in a second group; it ranges between +1 where all values of one group are higher than the other group, and −1 with the reverse relationship (two identical data sets would have a Cliff’s delta of 0). Both significance testing and power calculations were avoided following recommended practice (Altman et al. 2000; Cumming 2012); the Mann Whitney *U* statistic was used to calculate *P* values for *pro forma* reporting exclusively. To indicate estimate precision, 95% confidence intervals (95CI) were calculated using bootstrap methods and reported in text and/or as error bars (Cumming 2012).

## Data availability statement

The data and code that support the findings of this study, along with example video files, are available in the Zenodo repository with the identifier https://doi.org/10.5281/zenodo.495697.

## Author Contributions

*Conceptualization*: RN, ZPW and ACC; *Methodology*: RN, MR, DC and AE; *Software*: JCS and RN; *Investigation*: RN (SNAC, single CAFE), AE (group CAFE, respiration, lipid, weight, genetics, systematic review), MR (activity, climbing, individual CAFE, SNAC, genetics, systematic review), and YLT (systematic review); *Resources*: RN, JCS, DC, ZPW (instrumentation); *Data Analysis*: RN (SNAC, single CAFE, respiration), MR (SNAC, single CAFE), AE (group CAFE, lipid, weight), YLT (systematic review); *Writing – Original Draft*: MR, AE; *Writing – Revision*: AE and ACC; *Visualization*: MR, AE, RN and ACC; *Supervision*: ZPW and ACC; *Project Administration*: ACC; *Funding Acquisition*: ZPW and ACC.

## Acknowledgements

We thank Nicola Poh Si En for technical assistance with the single-fly CAFE, and Joses Ho for help plotting the systematic review results. We thank Jessica Edwards of Insight Editing London for editing of the manuscript.

## References

Al-Anzi, B. et al., 2009. Obesity-blocking neurons in Drosophila. Neuron, 63(3), pp.329–341.

Alekseyenko, O.V., Lee, C. & Kravitz, E.A., 2010. Targeted manipulation of serotonergic neurotransmission affects the escalation of aggression in adult male Drosophila melanogaster. PloS one, 5(5),p.e10806.

Altman, D. et al., 2000. Statistics with confidence: confidence interval and statistical guidelines. Bristol: BMJ Books.

Aso, Y. et al., 2014. The neuronal architecture of the mushroom body provides a logic for associative learning. eLife, 23(3), p.04577.

Baines, R.A. et al., 2001. Altered electrical properties in Drosophila neurons developing without synaptic transmission. The Journal of neuroscience: the o?cial journal of the Society for Neuroscience, 21(5), pp.1523–1531.

Baines, R.A. et al., 1999. Postsynaptic expression of tetanus toxin light chain blocks synaptogenesis in Drosophila. Current biology: CB, 9(21), pp.1267–1270.

Baker, K.D. & Thummel, C.S., 2007. Diabetic larvae and obese flies-emerging studies of metabolism in Drosophila. Cell metabolism, 6(4), pp.257–266.

Bartholomew, N.R. et al., 2015. Impaired climbing and fiight behaviour in Drosophila melanogaster following carbon dioxide anaesthesia. Scientific reports, 5, p.15298.

Burke, C.J. et al., 2012. Layered reward signalling through octopamine and dopamine in Drosophila. Nature, 492(7429), pp. 433–437.

Claridge-Chang, A. & Assam, P.N., 2016. Estimation statistics should replace significance testing. Nature methods, 13(2), pp. 108–109.

Clifi, N., 1993. Dominance statistics: Ordinal analyses to answer ordinal questions. Psychological bulletin, 114(3), p.494.

Cole, S.H. et al., 2005. Two functional but noncomplementing Drosophila tyrosine decarboxylase genes: distinct roles for neural tyramine and octopamine in female fertility. The Journal of biological chemistry, 280(15), pp.14948–14955.

Coulom, H. & Birman, S., 2004. Chronic exposure to rotenone models sporadic Parkinson’s disease in Drosophila melanogaster. The Journal of neuroscience: the official journal of the Society for Neuroscience, 24(48), pp.10993–10998.

Crocker, A. et al., 2010. Identification of a neural circuit that underlies the effects of octopamine on sleep:wake behavior. Neuron,

Crocker, A. & Sehgal, A., 2008. Octopamine regulates sleep in drosophila through protein kinase A-dependent mechanisms. The Journal of neuroscience: the official journal of the Society for Neuroscience, 28(38), pp.9377–9385.

Cumming, G., 2012. Understanding the new statistics effect sizes, confidence intervals, and meta-analysis, New York: Routledge.

Depetris-Chauvin, A. et al., 2011. Adult-specific electrical silencing of pacemaker neurons uncouples molecular clock from circadian outputs. Current biology: CB, 21(21), pp.1783–1793.

Friggi-Grelin, F. et al., 2003. Targeted gene expression in Drosophila dopaminergic cells using regulatory sequences from tyrosine hydroxylase. Journal of neurobiology, 54(4), pp.618–627.

Gasque, G. et al., 2013. Small molecule drug screening in Drosophila identifies the 5HT2A receptor as a feeding modulation target. Scientific reports, 3(10).

Harris-Warrick, R.M. & Marder, E., 1991. Modulation of neural networks for behavior. Annual review of neuroscience, 14, pp.39–57.

Inagaki, H.K. et al., 2012. Visualizing neuromodulation in vivo:TANGO-mapping of dopamine signaling reveals appetite control of sugar sensing. Cell, 148(3), pp.583–595.

Islam, R. et al., 2012. A neuroprotective role of the human uncoupling protein 2 (hUCP2) in a Drosophila Parkinson’s disease model. Neurobiology of disease, 46(1), pp.137–146.

Ja, W.W. et al., 2007. Prandiology of Drosophila and the CAFE assay. Proceedings of the National Academy of Sciences of the United States of America, 104(20), pp.8253–8256.

Jeong, Y.T. et al., 2016. Mechanosensory neurons control sweet sensing in Drosophila. Nature communications, 7, p.12872.

Kaun, K.R., Devineni, A.V. & Heberlein, U., 2012. Drosophila melanogaster as a model to study drug addiction. Human genetics, 131(6), pp.959–975.

Kim, W.J., Jan, L.Y. & Jan, Y.N., 2012. Contribution of visual and circadian neural circuits to memory for prolonged mating induced by rivals. Nature neuroscience, 15(6), pp.876–883.

Lee, K.-S. et al., 2004. Drosophila short neuropeptide F regulates food intake and body size. The Journal of biological chemistry, 279(49), pp.50781–50789.

Leibel, R.L., Rosenbaum, M. & Hirsch, J., 1995. Changes in energy expenditure resulting from altered body weight. The New England journal of medicine, 332(10), pp.621–628.

Li, H. et al., 2000. Ectopic G-protein expression in dopamine and serotonin neurons blocks cocaine sensitization in Drosophila melanogaster. Current biology: CB, 10(4), pp.211–214.

Liu, C. et al., 2012. A subset of dopamine neurons signals reward for odour memory in Drosophila. Nature, 488(7412), pp.512–516.

Loftus, T.M. et al., 2000. Reduced food intake and body weight in mice treated with fatty acid synthase inhibitors. Science, 288(5475), pp.2379–2381.

Mann, K., Gordon, M.D. & Scott, K., 2013. A Pair of Interneurons In?uences the Choice between Feeding and Locomotion in Drosophila. Neuron, 79(4), pp.754–765.

Marella, S., Mann, K. & Scott, K., 2012. Dopaminergic modulation of sucrose acceptance behavior in Drosophila. Neuron, 73(5), pp. 941–950.

Masek, P. & Keene, A.C., 2013. Drosophila fatty acid taste signals through the PLC pathway in sugar-sensing neurons. PLoS genetics, 9(9), p.e1003710.

McGuire, S.E., Mao, Z. & Davis, R.L., 2004. Spatiotemporal gene expression targeting with the TARGET and gene-switch systems in Drosophila. Science’s STKE: signal transduction knowledge environment, 2004(220), p.6.

Melcher, C., Bader, R. & Pankratz, M.J., 2007. Amino acids, taste circuits, and feeding behavior in Drosophila: towards understanding the psychology of feeding in flies and man. The Journal of endocrinology, 192(3), pp.467–472.

Navawongse, R. et al., 2016. Drosophila learn efficient paths to a food source. Neurobiology of learning and memory, 131, pp.176–181.

Neckameyer, W.S., 2010. A trophic role for serotonin in the development of a simple feeding circuit. Developmental neuroscience, 32(3), pp.217–237.

Obici, S. et al., 2002. Central administration of oleic acid inhibits glucose production and food intake. Diabetes, 51(2), pp.271–275.

Osterwalder, T. et al., 2001. A conditional tissue-specific transgene expression system using inducible GAL4. Proceedings of the National Academy of Sciences of the United States of America, 98(22), pp.12596–12601.

Owusu-Ansah, E. & Perrimon, N., 2014. Modeling metabolic homeostasis and nutrient sensing in Drosophila: implications for aging and metabolic diseases. Disease models & mechanisms, 7(3), pp.343–350.

Pandey, U.B. & Nichols, C.D., 2011. Human disease models in Drosophila melanogaster and the role of the fly in therapeutic drug discovery. Pharmacological reviews, 63(2), pp.411–436.

Pfa, D.W., Kieffer, B.L. & Swanson, L.W., 2008. Mechanisms for the regulation of state changes in the central nervous system: an introduction. Annals of the New York Academy of Sciences, 1129, pp.1–7.

Pool, A.-H. et al., 2014. Four GABAergic Interneurons Impose Feeding Restraint in Drosophila. Neuron, 83(1), pp.164–177.

Riemensperger, T. et al., 2013. A single dopamine pathway underlies progressive locomotor defficits in a Drosophila model of Parkinson disease. Cell reports, 5(4), pp.952–960.

Riemensperger, T. et al., 2011. Behavioral consequences of dopamine deficiency in the Drosophila central nervous system. Proceedings of the National Academy of Sciences of the United States of America, 108(2), pp.834–839.

Roeder, T., 2005. Tyramine and octopamine: ruling behavior and metabolism. Annual review of entomology, 50, pp.447–477.

Rogman, J.J., 2013. Ordinal Dominance Statistics (Orddom): an RProject for Statistical Computing Package to Compute Ordinal, Nonparametric Alternatives to Mean Comparison., Version 3.1. Available at: http://cran.r-project.org/.

Schoofs, A. et al., 2014. Selection of motor programs for suppressing food intake and inducing locomotion in the Drosophila brain. PLoS biology, 12(6).

Shaltiel-Karyo, R. et al., 2012. A novel, sensitive assay for behavioral defects in Parkinson’s disease model Drosophila. Parkinson’s disease, 697564(10), p.25.

Shen, P. & Cai, H.N., 2001. Drosophila neuropeptide F mediates integration of chemosensory stimulation and conditioning of the nervous system by food. Journal of neurobiology, 47(1), pp.16–25.

Shih, H.-W. et al., 2015. Parallel circuits control temperature preference in Drosophila during ageing. Nature communications, 6, p.7775.

Shimada-Niwa, Y. & Niwa, R., 2014. Serotonergic neurons respond to nutrients and regulate the timing of steroid hormone biosynthesis in Drosophila. Nature communications, 5(5778).

Vucetic, Z. & Reyes, T.M., 2010. Central dopaminergic circuitry controlling food intake and reward: implications for the regulation of obesity. Wiley interdisciplinary reviews. Systems biology and medicine, 2(5), pp.577–593.

Williams, M.J. et al., 2014. Obesity-linked homologues TfAP-2 and Twz establish meal frequency in Drosophila melanogaster. PLoS genetics, 10(9).

Wu, Q. et al., 2003. Developmental control of foraging and social behavior by the Drosophila neuropeptide Y-like system. Neuron, 39(1), pp.147–161.

Yang, Z. et al., 2015. Octopamine mediates starvation-induced hyperactivity in adult Drosophila. Proceedings of the National Academy of Sciences of the United States of America, 112(16), pp.|p5219– 5224.

Yildizoglu, T. et al., 2015. Estimating Information Processing in a Memory System: The Utility of Meta-analytic Methods for Genetics. PLoS genetics, 11(12), p.e1005718.

Zhang, L. et al., 2015. TMC-1 attenuates C. elegans development and sexual behaviour in a chemically defined food environment. Nature communications, 6, p.6345.

Zhang, T., Branch, A. & Shen, P., 2013. Octopamine-mediated circuit mechanism underlying controlled appetite for palatable food in Drosophila. Proceedings of the National Academy of Sciences of the United States of America, 110(38), pp.15431–15436.

Zhou, C. & Rao, Y., 2008. A subset of octopaminergic neurons are important for Drosophila aggression. Nature neuroscience, 11(9), pp.1059–1067.

Zhou, Q.Y. & Palmiter, R.D., 1995. Dopamine-defficient mice are severely hypoactive, adipsic, and aphagic. Cell, 83(7), pp.1197– 1209.

